# Tissue Inhibitor of Metalloproteinase 3 (TIMP3) mutations increase glycolytic activity and dysregulate glutamine metabolism in RPE cells

**DOI:** 10.1101/2024.01.05.574411

**Authors:** Allison Grenell, Charandeep Singh, Monisha Raju, Alyson Wolk, Sonal Dalvi, Geeng-Fu Jang, John S. Crabb, Courtney E. Hershberger, Kannan Manian, John W. Crabb, Ruchira Singh, Jianhai Du, Bela Anand-Apte

## Abstract

Mutations in Tissue Inhibitor of Metalloproteinases 3 (TIMP3) cause Sorsby’s Fundus Dystrophy (SFD), a dominantly inherited, rare form of macular degeneration that results in vision loss. TIMP3 is synthesized primarily by retinal pigment epithelial (RPE) cells, which constitute the outer blood-retinal barrier. Quantitative proteomics and RNAseq analysis on the choroid/RPE of mice expressing mutant TIMP3 identified a dysregulation in metabolic processes. We examined the effects of mutant TIMP3 on RPE metabolism using human ARPE-19 cells expressing mutant S179C TIMP3 and patient-derived induced pluripotent stem cell-derived RPE (iRPE) carrying the S204C TIMP3 mutation. Stable isotope tracing experiments demonstrated enhanced glucose utilization and glycolytic activity in mutant RPE concomitantly with altered glutamine utilization. This study provides important information on the dysregulation of the metabolome of RPE cells in SFD and implicates a potential commonality with other retinal degenerative diseases, emphasizing RPE cellular metabolism as a therapeutic target.

## Introduction

Sorsby’s Fundus Dystrophy (SFD) is a rare autosomal dominant form of macular degeneration which results in progressive vision loss starting in the 3rd to 4th decade of life. Mutations in the Tissue Inhibitor of Metalloproteinases 3 (TIMP3) gene cause the disease with 100% penetrance. TIMP3 is an extracellular inhibitor of matrix metalloproteinases (MMPs) which are involved in the degradation of extracellular matrix (ECM) proteins such as collagen and elastin. TIMP3 is synthesized by the retinal pigment epithelium (RPE) and endothelial cells of the choriocapillaris in the retina. The pathological features of SFD bear a striking similarity to the more common age-related macular degeneration (AMD) with defects in Bruch’s membrane (BrM), subretinal deposits (“drusen”) and choroidal neovascularization (CNV)^1–4^. Drusen in both AMD and SFD contain TIMP3, leading some to speculate that they share an underlying etiology^4,5^.

Recently, metabolic dysfunction has emerged as an important contributor to AMD ^6–8^ as well as other forms of retinal degeneration including retinitis pigmentosa^9^, diabetic retinopathy^10,11^, retinopathy of prematurity^12,13^, and macular telangiectasia (MacTel)^14–16^. Mutations in genes associated with metabolism (glycolysis^17,18^, fatty acid beta oxidation^19^, tri-carboxylic acid cycle (TCA)^18^, and nucleotide synthesis^20,21^) have been shown to cause retinal degeneration. In addition, some system-wide metabolic diseases, are known to negatively impact the retina such as diabetic retinopathy (DR), a vision threatening complication of diabetes mellitus^20^.

Metabolic perturbations such as hypoxia-induced metabolic stress, specifically in the RPE, cause photoreceptor degeneration in mice^22^. RPE cells constitute the outer blood retinal barrier and are essential for photoreceptor health and function. One important RPE function is the transport and production of vital nutrients, such as glucose, to the retina. Glucose is needed to support the heavy reliance of photoreceptors on aerobic glycolysis, where glucose is metabolized to lactate in the cytosol to generate ATP^23,24^. Under physiological conditions, RPE spares glucose for photoreceptors by utilizing alternative fuels including proline^25,26^, fatty acids^27^, glutamine^28^ and lactate^29^. The disruption of metabolic coupling between RPE and photoreceptors result in the degeneration of photoreceptors^9^. Additionally, altered mitochondrial metabolism in the RPE of human AMD donor eyes underscores the importance of RPE metabolism in sustaining the retinal metabolic ecosystem^30^.

We examined metabolic changes in the RPE expressing mutant TIMP3 that might contribute to the pathogenesis of SFD. Quantitative proteomics and transcriptomics experimental analysis on RPE/posterior eye cups of SFD mice carrying the TIMP3 S179C variant identified dysregulated glycolysis pathways. Using stable isotope labeling, we directly analyzed the effects of mutant TIMP3 on RPE metabolism using two *in vitro* models of SFD: ARPE-19 S179C TIMP3 and induced pluripotent stem cell (iPSC) derived RPE from patients with the S204C TIMP3 mutation (iRPE). Our data supports the hypothesis that TIMP3 mutations leads to increased glycolytic activity and glucose contribution to the TCA cycle in RPE cells.

## Results

### Transcriptomics and proteomics of RPE in mice expressing TIMP3S179C reveal abnormalities in metabolic processing

To understand the molecular and cellular processes involved in SFD, quantitative proteomics and RNAseq were performed on RPE and posterior eye cups of mice that globally expressed the S179C variant of TIMP3 and compared with wild-type littermates. PANTHER Gene Ontology (GO) analysis of 298 proteins that were differentially expressed in the RPE of mutant mice, (adjusted p < 0.05) were most significantly overrepresented in the ‘metabolite interconversion enzyme’ protein class (GO: PC00262) comprising 17% of all differentially expressed proteins (p < 0.05) (**Figure 1A, Supporting Information 1**). The most overrepresented cellular process was the ‘cellular metabolic process’ (GO:0044237) with 18% of differentially expressed proteins falling into this category (p < 0.05) (**Figure 1B, Supporting Information 1**).

**Figure 1:**
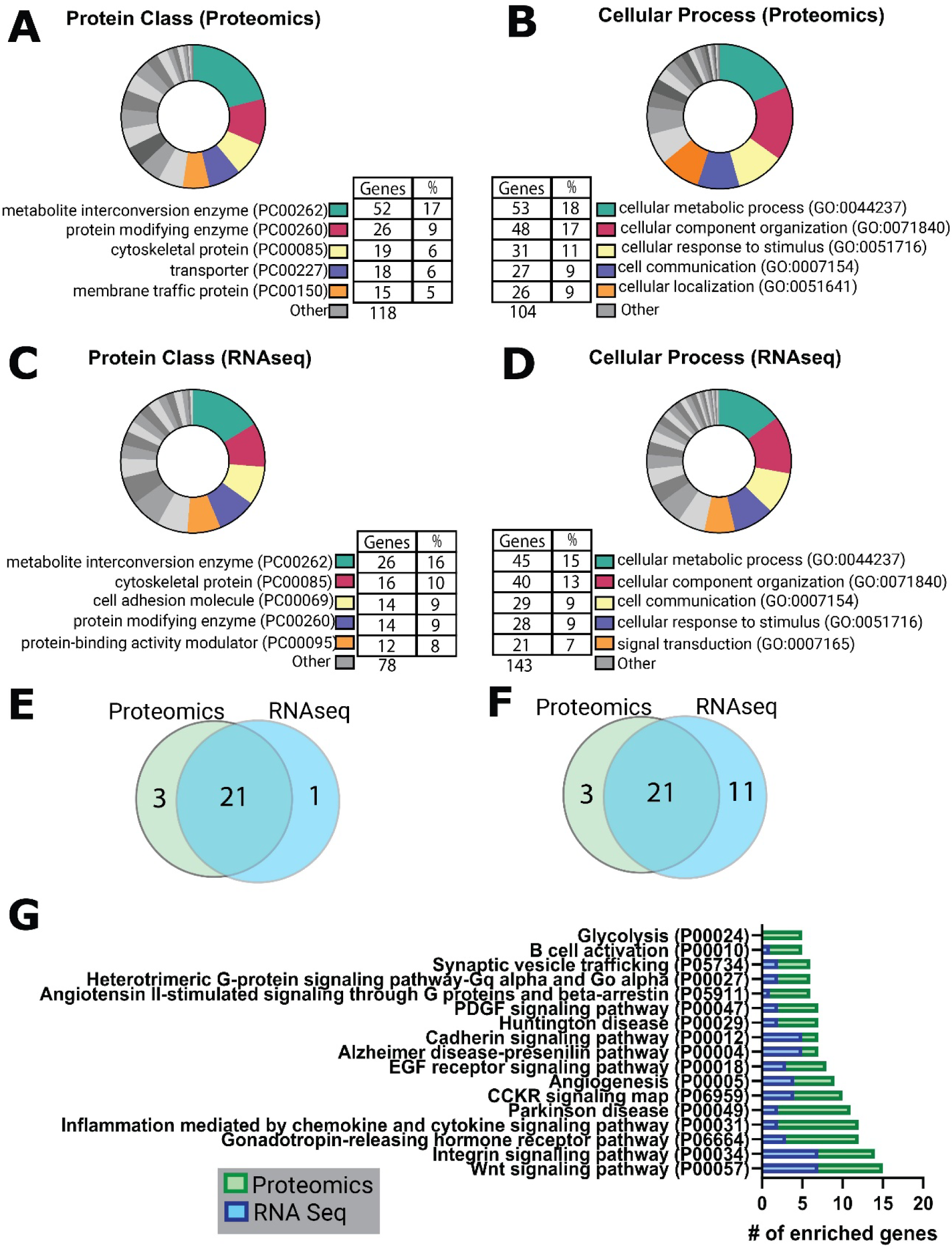
Panther Gene Ontology analysis of proteins and transcripts found to be significantly altered in S179C TIMP3 mice RPE/eyecups when compared to WT controls. Proteomics and RNA seq was conducted on TIMP3 S179C RPE/eye cups and WT controls (n=8). Panther Gene Ontology (GO) analysis was carried out on proteins and transcripts found to be significantly altered (adjusted p value < 0.05) between the two groups. A, Top 5 differential protein classes identified from proteomics along with the number of genes altered in each pathway (“genes”) and percent of genes hit against total # genes (“%”). B, Top 5 differential cellular process identified from proteomics. C, Top 5 differential protein classes identified from RNAseq D, Top 5 differential cellular processes identified from RNAseq. E, Comparison of proteomics and RNAseq results identified an overlapping list of dysregulated protein classes and F, cellular processes. G, pathways containing the highest number of dysregulated genes.

Panther GO analysis was also performed on 212 differentially expressed transcripts discovered by RNAseq (adjusted P<0.05) **(Supporting Information 2).** Like the proteomics analysis, the differentially expressed genes were most overrepresented in ‘metabolite interconversion enzyme’ and ‘cellular metabolic process’ of the GO protein classes and cellular processes, respectively (p < 0.05) **(Figure 1C, D)**. Differential expression of both genes and proteins involved in the metabolic pathways was a strong indication of this pathway being regulated with mutant TIMP3 in RPE. Between the two datasets, differential expression was found in 25 total protein classes and of those 21 (∼84%) were identified in both proteomics and RNAseq analyses (p < 0.05) **(Figure 1E).** Likewise, 35 cellular processes were enriched with 21 (∼60%) shared (p < 0.05) **(Figure 1F).** Together, these data this strongly support the hypothesis that TIMP3S179C mutation perturbs the metabolism of the RPE.

Finally, pathway analysis using Panther GO were performed on all differentially expressed genes in the RNAseq and proteomics data sets. (**Figure 1G**). Our data confirmed the role of angiogenesis which is known to be largely dysregulated in SFD (p < 0.05). Five differentially expressed proteins were found to be significantly overrepresented in the glycolysis pathway, by Panther GO **(Figure 6A, Supporting Information 3)**. Notably bisphosphoglycerate mutase (BPMG) protein levels were increased. BPGM, facilitates the conversion of the glycolytic intermediate 1,3 bisphosphoglycerate to 2,3 bisphosphoglycerate and regulates glycolytic intermediates and serine synthesis^31^. Concurrently, Hexokinase 1 (HK1), Alpha enolase (ENO1), gamma-enolase (ENO2), and pyruvate kinase (PK) were found to be down-regulated. Previous studies in tumors demonstrated that knocking down HK1, responsible for the rate-limiting step in glycolysis, led to dysregulated energy metabolism with increased glycolysis. ENO1 and ENO2 catalyze the conversion of 2-phosphoglycerate to phosphoenolpyruvate while PK catalyzes the synthesis of pyruvate from phosphoenolpyruvate pyruvate.

Other pathways known to regulate metabolism, albeit indirectly were also identified. Wnt signaling^32^ and integrin signaling^33^, is well known for regulate or be regulated by key metabolic enzymes and transcription factors.

### TIMP3^S179C^ ARPE-19 cells have increased glycolytic activity

Uniformly labeled [^13^C_6_] glucose isotopic tracing was performed on ARPE19 cells^34^, an immortalized human RPE cell line, that was engineered to expressed the S179C TIMP3 variant^35^, to directly study metabolism and infer glycolytic activity. Comparisons were made between RPE cells expressing mutant RPE and cells engineered to express equivalent amounts of wild-type (WT++) TIMP3 to directly study metabolism and infer glycolytic activity (**Figure 2A**). ARPE-19 cells were cultured using a method previously shown to increase epithelial cell morphology, cytoskeletal organization and RPE-related functions^36^. All non-labeled glucose in the cell culture media was replaced by 10mM [^13^C_6_] glucose. Cells were incubated with the labeled media for 24 hours to ensure that isotopic steady state was reached before metabolite extraction. To study glycolysis specifically, relative levels of intracellular lactate **(Figure 2B)** and pyruvate **(Figure 2C)** were evaluated compared to controls. An increase in intracellular pyruvate and lactate in TIMP3 S179C mutant RPE suggests increased glycolytic activity.

**Figure 2:**
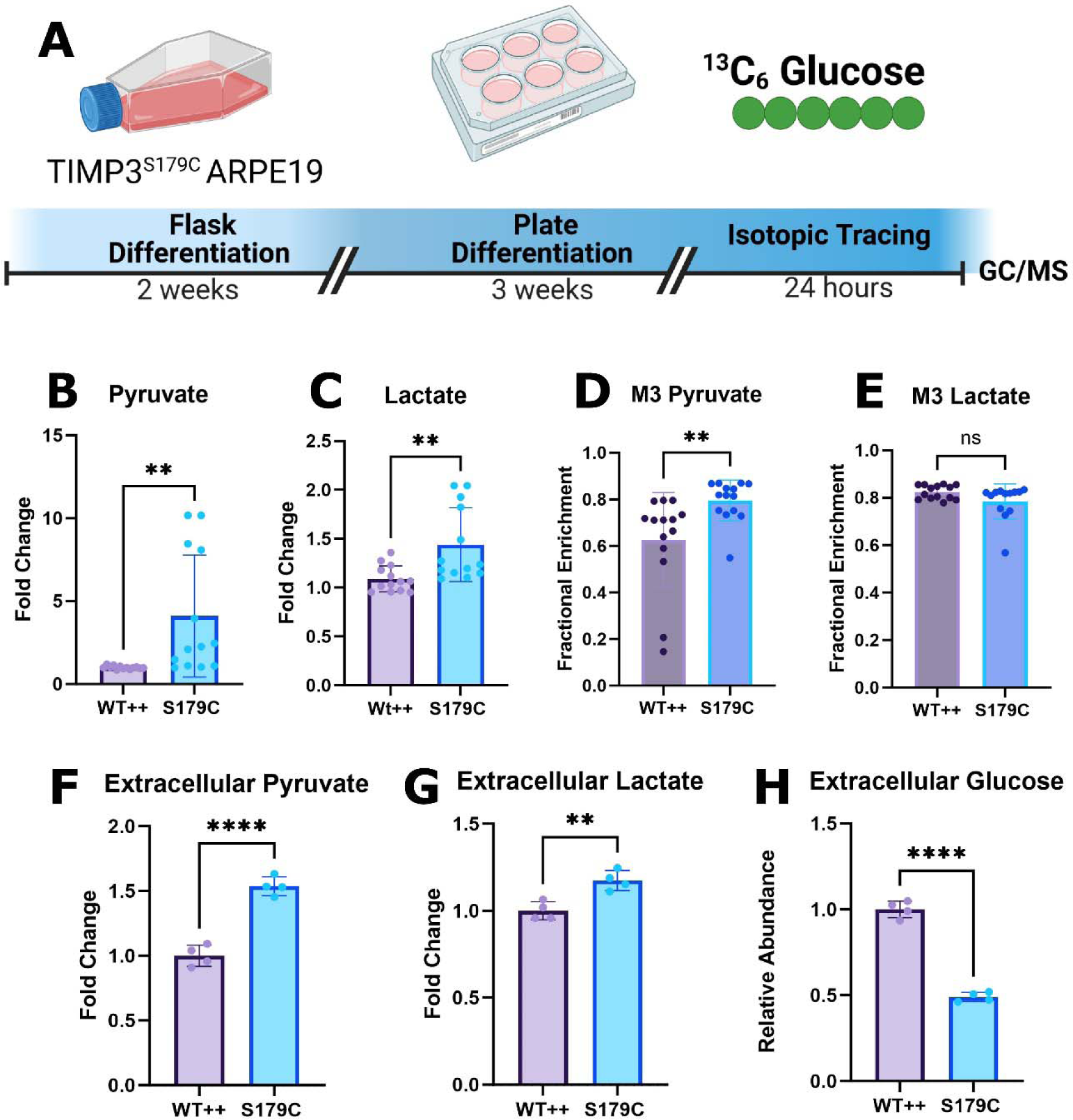
TIMP3S179C ARPE-19 cells have increased glycolytic activity. A, ARPE19 cells overexpressing S179C TIMP3 (S179C) or wild type TIMP3 (WT++) were fully differentiated before steady state [^13^C_6_] Glucose isotopic tracing using GC/MS was conducted for 24 hours. B, C, Total intracellular pyruvate and lactate levels were increased in S179C cells (n=13) D, M3 Pyruvate enrichment was increased in S179C cells while E, M3 Lactate remained unchanged. (n=14) F, G, The conditioned media was evaluated for extracellular pyruvate and lactate. Both were increased indicating increased production (n=4). H: Glucose levels were evaluated which were reduced by ∼50% in the S179C ARPE-19 cells indicating increased glucose utilization. (n=4). Statistics performed were 2 tailed unpaired t-tests. *p < 0.05, **p < 0.01, *** p < 0.001, **** p < 0.0001. Experimental schematic was generated using BioRender. Graphs were made with GraphPad Prism

Fully labeled M6 glucose (all 6 carbons labeled) is broken down via glycolysis to generate fully labeled M3 (3 carbons labeled) pyruvate. Under steady state conditions, an average of 64% of pyruvate was M3 labeled in WT APRE19 while 79% was M3 labeled in TIMP3 S179C cells **(Figure 2D)**. Increased M3 Pyruvate enrichment supports increased glucose utilization and glycolytic flux in TIMP3 S179C cells. Pyruvate can also undergo aerobic glycolysis to produce M3 lactate (**Figure 2E**). After 18 hours, the spent media was collected for extracellular analysis (**Figure F, G**). Consistent with results obtained from the intracellular analysis, extracellular pyruvate and lactate were elevated in S179C ARPE-19 media, further supporting an increase in glycolytic activity.

Finally, glucose levels were measured in conditioned medium of the cells after 18 hours to evaluate glucose utilization. Glucose levels were significantly decreased in TIMP3 S179C cells, confirming increased glucose utilization (**Figure 2H**). Collectively, these data suggest a metabolic dysregulation in RPE cells expressing the TIMP3 S179C mutation. More specifically, mutant TIMP3 results in increased glucose utilization and glycolytic flux in the RPE cells.

### SFD iRPE display dysregulated central carbon metabolism

To confirm that these findings were not limited to the immortalized ARPE-19 cell line, we performed similar studies on an human induced pluripotent stem cell (iPSC) model of SFD^37^ (iRPE) to validate our results and further characterize RPE central carbon metabolism. This model was previously shown to recapitulate hallmarks of healthy RPE and as well as AMD RPE^38,39^. Cells were originally collected through skin biopsies from patients with the TIMP3 S204C variant of SFD. Patient fibroblasts were dedifferentiated to iPSC cells and then re-differentiated into iRPE cells (**Figure 3A**). Cells with the mutation corrected by CRISPR-Cas9 were used as a control.

**Figure 3:**
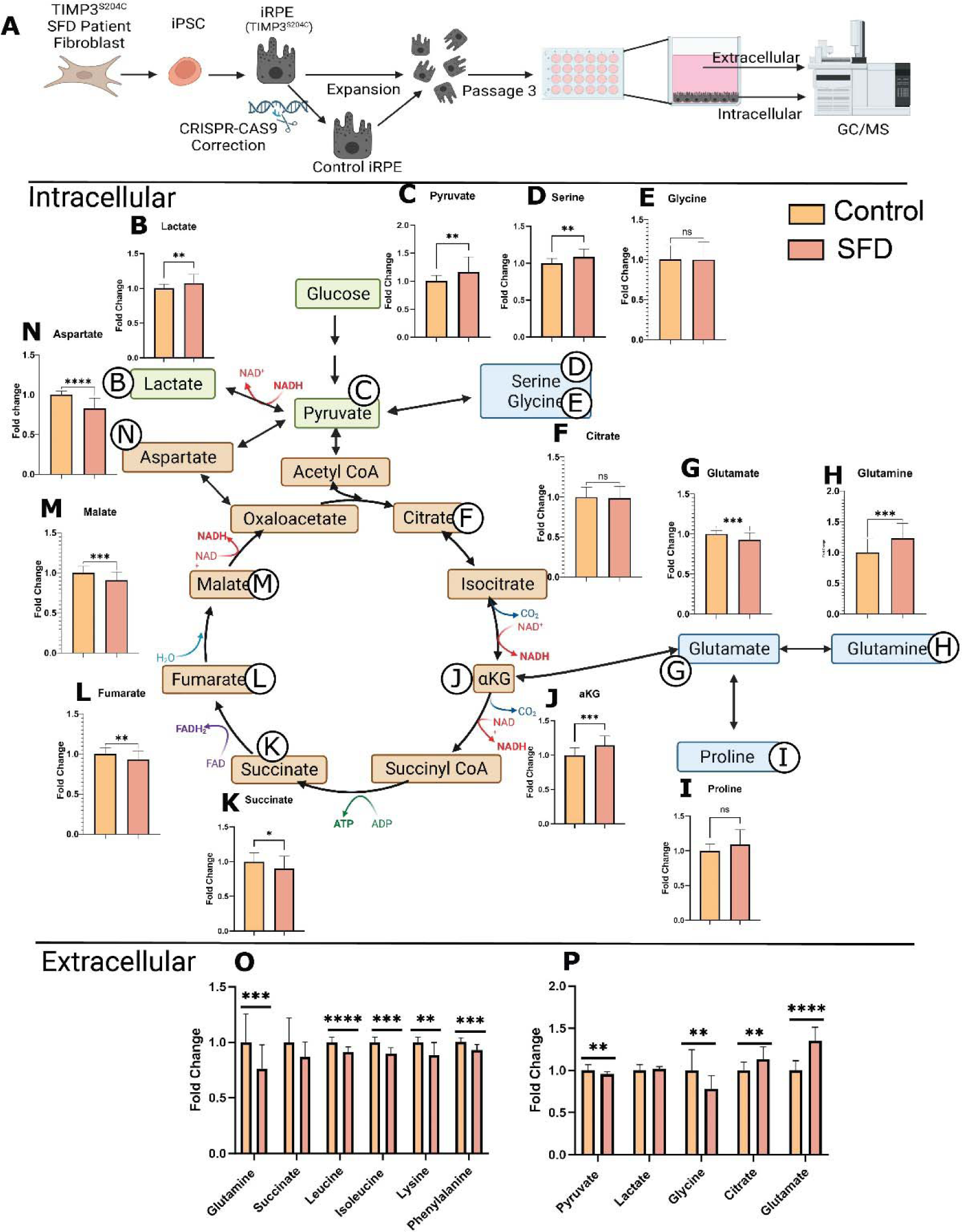
SFD iRPE cells display dysregulated central carbon metabolism. A, Fibroblasts from SFD patients with the S204C mutation were reprogrammed into iPSCs which were then differentiated into iPSC-RPE cells (iRPE). Fibroblasts from SFD patients with the S204C mutation were de-differentiated before being re-differentiated into RPE cells (iRPE). Cells were expanded until passage 3 and then cultured for a minimum 30 days. Relative intracellular and extracellular (from conditioned media) metabolites were quantified with GC/MS. B-N, Relative abundance of total metabolite levels was compared between SFD iRPE (SFD) and Crispr corrected controls (Control). Extracellular metabolites in conditioned media were quantified after 24 hours. O: metabolites that were decreased after 24 hours compared to time 0. P. metabolites that were increased after 24 hours compared with time 0. Statistics performed were two-tailed unpaired t-tests. *p < 0.05, **p < 0.01, *** p < 0.001, **** p < 0.0001. TCA cycle schematic was generated using BioRender. Graphs were made with GraphPad Prism (n=26)

As with ARPE-19, we found evidence of dysregulated central glucose metabolism in SFD iRPE cells. Consistently, there were increased levels of intracellular lactate (**Figure 3B**) and pyruvate (**Figure 3C**) suggesting increased glycolytic activity. Serine and glycine (**Figure 3D, E**) are metabolites generated when the glycolytic intermediate 3-phosphoglycerate (3-PG) is diverted by a 3-step enzymatic reaction that generates serine which can then be converted directly to glycine to feed carbon into one-carbon pathway. Serine levels **(Figure 3D)** but not glycine levels **(Figure 3E)** were found to be significantly increased indicating increased glycolysis with higher production of serine with no change in serine utilization downstream into the one-carbon pathway.

iRPE had increased intracellular alpha ketoglutarate (αKG) **(Figure 3J)** and glutamine **(Figure 3H).** αKG is primarily generated within the TCA cycle from citrate. Glutamine can also be generated from αKG through cataplerosis in which TCA intermediates exit the cycle and are redirected for biosynthesis. Intermediates that follow αKG in the TCA cycle were decreased in SFD iRPE, including succinate **(Figure 3K)**, fumarate **(Figure 3L),** malate **(Figure 3M)**, and aspartate **(Figure 3N)**. Glutamine accumulation with concurrent TCA intermediate depletion implies increased cataplerosis in SFD iRPE cells.

Semi-targeted extracellular metabolomics was conducted on the media to evaluate nutrient/metabolite uptake and release in SFD iRPE cells. After 24 hours, the relative total amount of metabolites (sum of all isotopologues) were compared between SFD iRPE and Crispr-corrected controls. **Figure 3O**, depicts metabolites that were utilized by the cells (present in the original media composition at time 0 and were decreased after 24 hours). There was increased utilization of glutamine, leucine, isoleucine, lysine and phenylalanine by SFD iRPE cells. **Figure 3P**, includes metabolites released by RPE cells (If levels were increased after 24 hours compared with time 0). There was a decrease in release of extracellular pyruvate and glycine and an increase in extracellular citrate and glutamate in SFD iRPE cells.

### SFD iRPE have increased glucose contribution to the TCA cycle and increased cataplerosis to glutamine

Since metabolite pool size alone is insufficient to deduce pathway activity or nutrient contribution, we used stable isotope tracing with [^13^C_6_] glucose to further characterize central carbon metabolism. When [^13^C_6_] glucose undergoes glycolysis, M3 lactate and M3 pyruvate are produced. Although intracellular M3 pyruvate enrichment was unchanged, Lactate enrichment was increased in SFD iRPE cells compared with the Crispr corrected control (**Figure 4 A, B).** This is consistent with increased glycolytic flux in SFD iRPE cells relative to control. Uniformly labeled [^13^C_6_] glucose produces M2, M4 and M6 citrate during the first, second, and third round of the TCA cycle respectively while M2 and M4 αKG, fumarate, malate, and aspartate are generated through the first and second round of the TCA cycle respectively. Increased M6 citrate indicates a slight increase in glucose contribution to the TCA cycle via Acetyl CoA (**Figure 4C**). This is supported by a concurrent increase in M4 fumarate, malate and aspartate enrichment (**Figure 4G-I**), suggesting increased TCA cycle activity. Pyruvate can alternatively enter the TCA cycle via pyruvate carboxylase (PC) which generates M3 aspartate, malate and fumarate (**Supporting Information 4**). No significant changes of these isotopologues were found, indicating that there is no change in pyruvate anaplerosis.

**Figure 4:**
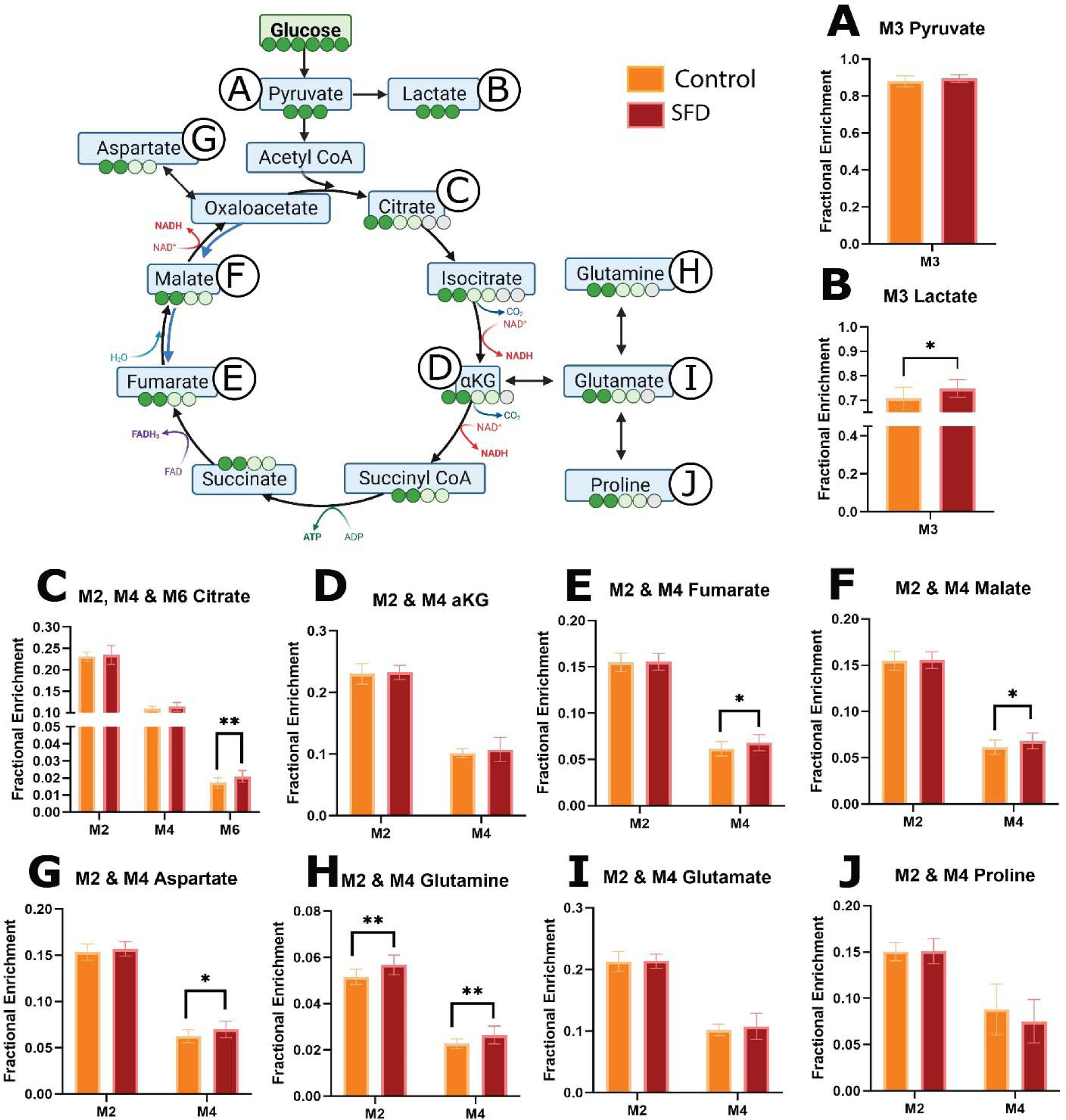
Stable isotope tracing using [^13^C_6_] glucose in SFD iRPE cells compared with controls. SFD iRPE cells have increased glycolytic activity, glucose contribution to the TCA cycle, and glutamine cataplerosis. Steady state isotopic tracing using [^13^C_6_] glucose was conducted, fractional enrichment was calculated and displayed on the Y axis. Isotopologues generated through glycolysis and TCA cycle are displayed on the X axis. A, B, When [^13^C_6_] glucose is broken down via glycolysis, M3 pyruvate and lactate are generated. C-I, M2 and M4 TCA cycle metabolites are generated during the first and second round of the TCA cycle respectively when pyruvate enters through Acetyl CoA. M2 and M4 glutamate, glutamine and proline are generated when cataplerosis occurs, and TCA intermediates are depleted. Statistics performed were 2 tailed unpaired t-tests. *p < 0.05, **p < 0.01, *** p < 0.001, **** p < 0.0001. TCA pathway schematic was generated using BioRender. Graphs were made with GraphPad Prism (n=13)

One of the main cataplerotic pathways from the TCA cycle involves the conversion of αKG to glutamate then glutamine. Increased M2 and M4 glutamine enrichment in SFD iRPE implies that glucose is being used to generate glutamine by leaving the TCA cycle. This is further supported by an increase in M5 glutamine and M5 proline **(Supplemental Data 4l)** which are generated from fully labeled citrate. In total, this demonstrates that SFD iRPE cells display increased TCA cycle activity and cataplerosis. This is likely contributing to the intracellular glutamine accumulation. It remains unclear why SFD iRPE cells would redirect nutrients allocated for energy production towards glutamine biosynthesis and how this additional glutamine will be utilized.

### SFD iRPE has no change in glutamine contribution to the TCA cycle and impaired malic enzyme activity

We demonstrated increased glutamine cataplerosis, or the removal of TCA intermediates, in SFD iRPE cells, however, anaplerosis, or the replenishment of TCA cycle intermediates, is a process that happens in parallel. Anaplerosis can be an important fate for intracellular glutamine. It involves its integration into the TCA cycle through αKG. Furthermore, increased anaplerosis has previously been identified in another model of retinal degeneration^40^. Therefore, glutamine metabolism was probed directly using uniformly labeled [^13^C_5_] glutamine isotopic tracing (**Figure 5A, Supporting Information 5**). SFD iRPE and Crispr corrected controls cells were incubated for 24 hours with media that contained labeled glutamine. Relative metabolite levels between the mutant and control cells were determined via GC/MS and mass isotopomer distributions (MIDs) were calculated after correction for natural isotopic abundance.

**Figure 5:**
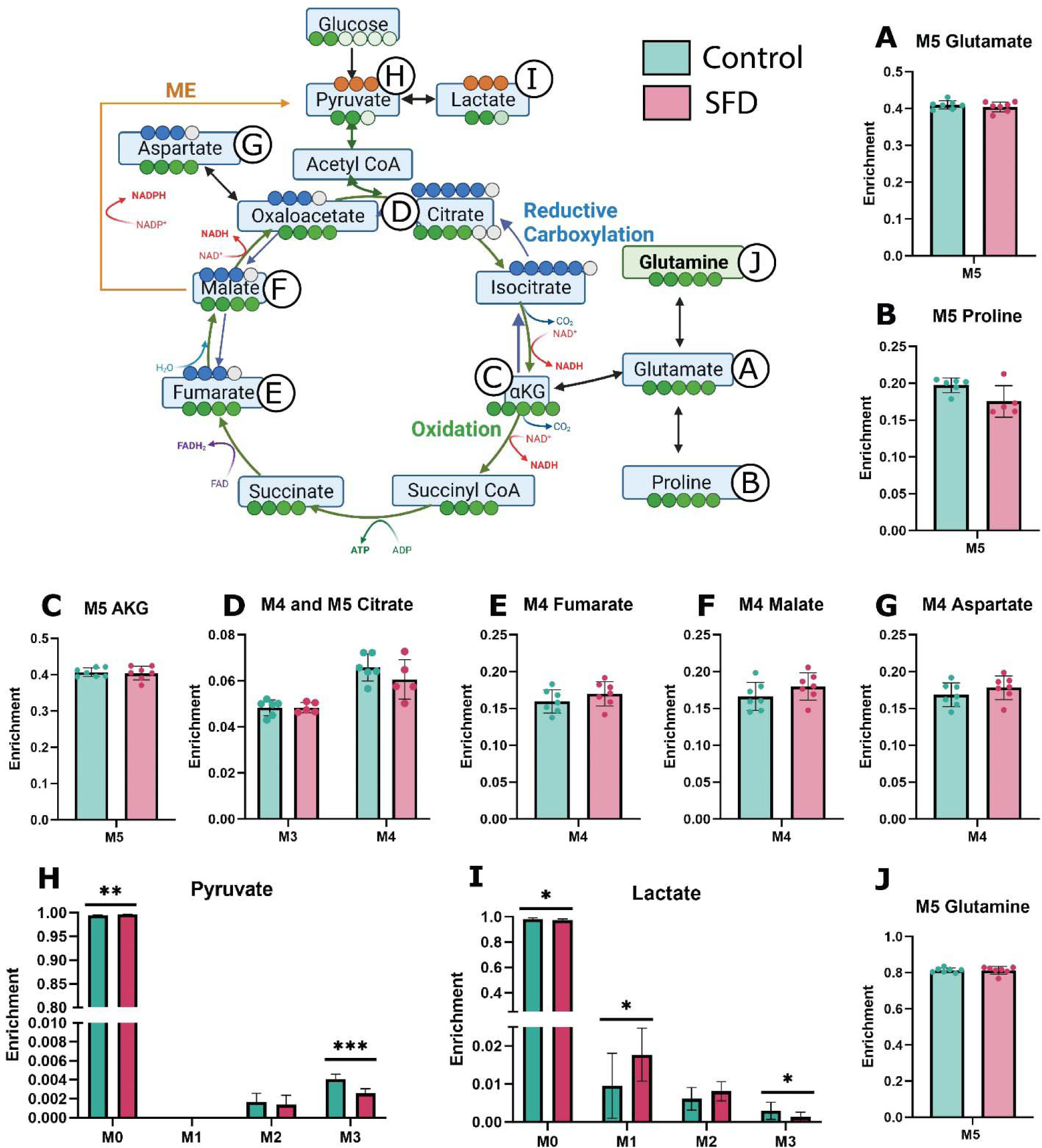
Stable isotope tracing using [^13^C_5_] glutamine in SFD iRPE cells compared with controls. Steady state isotopic tracing with [^13^C_5_] Glutamine was conducted, and fractional enrichment was calculated and is displayed on the Y axis. M5 glutamine is metabolized to M5 glutamate (A) which converted to either M5 proline (B) or M5 αkg (C). M4 fumarate or M5 citrate (D) or M4 fumarate (E) are produced from reductive carboxylation or oxidation of αKG respectively. Forward flow of the TCA cycle results in M4 Malate (F), M4 Aspartate (G), and M4 Citrate (D). Carbons can exit the TCA cycle to generate M2 Pyruvate (H) and M2 Lactate (I) via Acetyl CoA or M3 Pyruvate and M3 Lactate via malic enzyme. (J) M5 glutamine represents glutamine taken up from the media. Individual data points represent biological replicates. Two tailed unpaired t-test was used to evaluate statistical significance. *p < 0.05, **p < 0.01, *** p < 0.001, **** p < 0.0001. Experimental schematic was generated using BioRender. Graphs were made with GraphPad Prism. (n=7)

After M5 glutamine can be converted to M5 glutamate (**Figure 5B**). M5 glutamate enrichment was similar in control and SFD iRPE indicating that glutamine entry and conversion to glutamate was unchanged in SFD iRPE.

When M5 glutamine is utilized by the TCA cycle, it generates M5 αKG along with M4 intermediates such as fumarate, malate, and aspartate. We found no change in TCA cycle M4 intermediate metabolite enrichment. Consistent with this, we found no change in M2 pyruvate and lactate which is formed when M4 intermediates, generated from glutamine, exit the TCA cycle. This indicates that glutamine anaplerosis remained unchanged in SFD iRPE **(Figure 5D-H).**

We observed a significant decrease in M3 lactate and pyruvate isotopic enrichment in SFD iRPE cells. M3 pyruvate and lactate is generated from uniformly labeled glutamine via mitochondrial malic enzyme, indicating potential inhibition of malic enzyme (ME) activity in iRPE cells (**Figure I-J**). This is consistent with a previous study using SFD iRPE cells suggesting decreased ME activity^2^. ME plays a pivotal role in converting malate to pyruvate while producing NADPH, a crucial factor for energy metabolism and maintaining cellular redox balance. Malic enzyme can also catalyze the reverse reaction of pyruvate to malate making it relevant for anaplerosis. It important to note that the amount of pyruvate and lactate generated from malic enzyme, even under control conditions, is relatively low at 0.5%. The biological relevance of this change will need to be confirmed. Malic enzyme generates NADPH, which plays a critical role in redox balance. We have previously shown that SFD iRPE cells have high oxidative stress at baseline and increased susceptibility to oxidative stress damage^2^. This together with our current data suggests that there is a potential loss of redox balance in SFD iRPE cells. Further research is required to ascertain the significance of ME activity and its role in the generation of NADPH within the RPE during healthy and diseased conditions.

## Discussion

Our data showing increased glycolytic activity and glucose consumption in ARPE-19 cells expressing mutant TIMP3 and SFD patient derived iRPE cells supports the hypothesis that TIMP3 mutations leads to increased glycolytic activity and glucose contribution to the TCA cycle in RPE. Increased glycolysis in the RPE has been shown to perturb retinal metabolism in retinitis pigmentosa^9^. While SFD and retinitis pigmentosa are caused by mutations in unique genes, our data suggests the possibility that altered glucose metabolism could be a common feature of these two retinal degenerative diseases and possibly others. RPE central carbon metabolism could be an attractive therapeutic target that could be used for both conditions.

The abnormalities in glycolysis in TIMP3 mutant RPE cells was also corroborated by proteomics. Panther GO analysis identified five dysregulated enzymes involved in glycolysis however no dysregulated glycolytic enzymes were identified by RNAseq. Glycolytic flux is regulated by four metabolic enzymes, glucose transporters (GLUTs), hexokinase (HK), phosphofructokinase (PFK), and lactate exporters (MCTs). Regulation of glycolytic flux occurs primarily at the protein production post-translational level, which includes allosteric regulation by ATP and pyruvate^41^. Additionally, the expression of glycolytic genes poorly correlate with the amounts of enzymes and downstream flux control^41,42^. For example, in hypoxia-induced glycolysis, the ATP/AMP ratio is decreased, which activates PFK^43^resulting in the production of F-1,6 bisphosphate, which further increases glycolytic flux. This process does not necessarily require a change in gene expression as flux is modulated by allosteric activation or inhibition of enzymes involved.

Glutamine was found to be to be the most altered metabolite intracellularly and extracellularly. Intracellular levels of glutamine were increased while extracellular glutamine was decreased in TIMP3 mutant RPE cells. Furthermore, [^13^C_6_] glucose isotopic tracing found that SFD iRPE cells have an increased propensity to synthesize glutamine from glucose. However, when glutamine metabolism was probed specifically with [^13^C_5_] glutamine isotopic tracing, there was no change in flux to intracellular glutamate or anaplerosis, however there was decreased malic enzyme (ME) activity. Although ME is a minor pathway for glutamine metabolism in RPE cells^28^, it is an important enzyme for NADPH production and crucial for the maintenance of redox balance. The relevance of ME activity in maintaining RPE redox homeostasis remains unknown and warrants future investigation.

Extracellularly, there was reduced glutamine and elevated glutamate in the conditioned media after 24 hours in culture, suggesting increased metabolism of glutamine to glutamate in SFD iRPE cells. Surprisingly, the intracellular glutamine levels were decreased in mutant RPE cells. In addition, intracellular glutamine-derived glutamate enrichment was unchanged in SFD iRPE cells. Similarly, we observed that pyruvate, which was increased intracellularly, was decreased extracellularly. Our results seem to refute the general dogma that extracellular metabolites, usually reflect RPE intracellular metabolism. While the reasons for the discrepancy between intracellular and extracellular metabolites remain obscure, there are several possibilities that might be worthy of exploration in the future.

One possibility is that there may be inefficient metabolite transport. Our proteomics data identified two differentially expressed glutamate transporters in mouse SFD RPE when compared with controls: excitatory amino acid transporter 1 (EAAT1) and mitochondrial glutamate transporter 1 (SLC25a22). If mutant RPE cells had increased export or decreased import of glutamine from the cell, this would potentially result in a decrease in intracellular and an increase in secreted extracellular levels of glutamate.

Alternatively, another explanation could be the presence of extracellular metabolism. TIMP3 is an extracellular protein and TIMP3 mutations result drastic changes to the extracellular environment, especially the ECM. While we typically think of metabolism as an intracellular process, extracellular metabolism occurs in other systems such as bacteria, plants, and even cancer cells. Extracellular space has been identified as an important compartment for malignant energetic catalysis in cancer^44^. Furthermore, it has been reported that extracellular vesicles (EVs) can contain active metabolic enzymes that are involved in glycolytic, nucleotide, nucleoside, and xenobiotic metabolism^45–47^. EVs are known to be released from RPE cells in response to stressors^48^ and exosome markers have been found in Bruch’s membrane^49^. Additionally, glycolytic enzymes are among the most abundant enzymes associated with EVs. Our results could be explained by decreased extracellular glycolysis concurrently with increased extracellular glutamine to glutamate conversion. If extracellular RPE metabolism were occurring, this would completely change our current understand of the retinal metabolic ecosystem.

It remains unclear why SFD iRPE cells increasingly use glucose to synthesize glutamine and how the glutamine is utilized. Since our results suggests that mutant RPE do not display increased glutamine anaplerosis or proline synthesis, it is possible that the glutamine may be utilized for the biosynthesis of amino acids, nucleotides, and other macromolecules. Thickened Bruch’s membrane is a common pathological feature in patients with SFD ^2,3,50^. It is possible that this is due to increased ECM biosynthesis because of increased glutamine. We have previously reported increased deposition of the extracellular glycosaminoglycan, Hyaluronan (HA), in SFD RPE cells and AMD patient samples^50^. HA is synthesized via the hexosamine biosynthetic pathway (HBP) which branches off from glycolysis. Glutamine: fructose-6-phosphate amido transferases (GFAT) are enzymes that utilize glutamine to catalyze the first rate-limiting step of the pathway. An intriguing hypothesis is that RPE cells utilize glutamine to support increased HBP activity (**Figure 6B**). Alternatively, increased HA production could be a byproduct of glycolytic spillover from glycolytic intermediate accumulation. Regardless, either scenario would require glutamine utilization and could explain why SFD iRPE cells are generating more glucose-derived glutamine.

**Figure 6:**
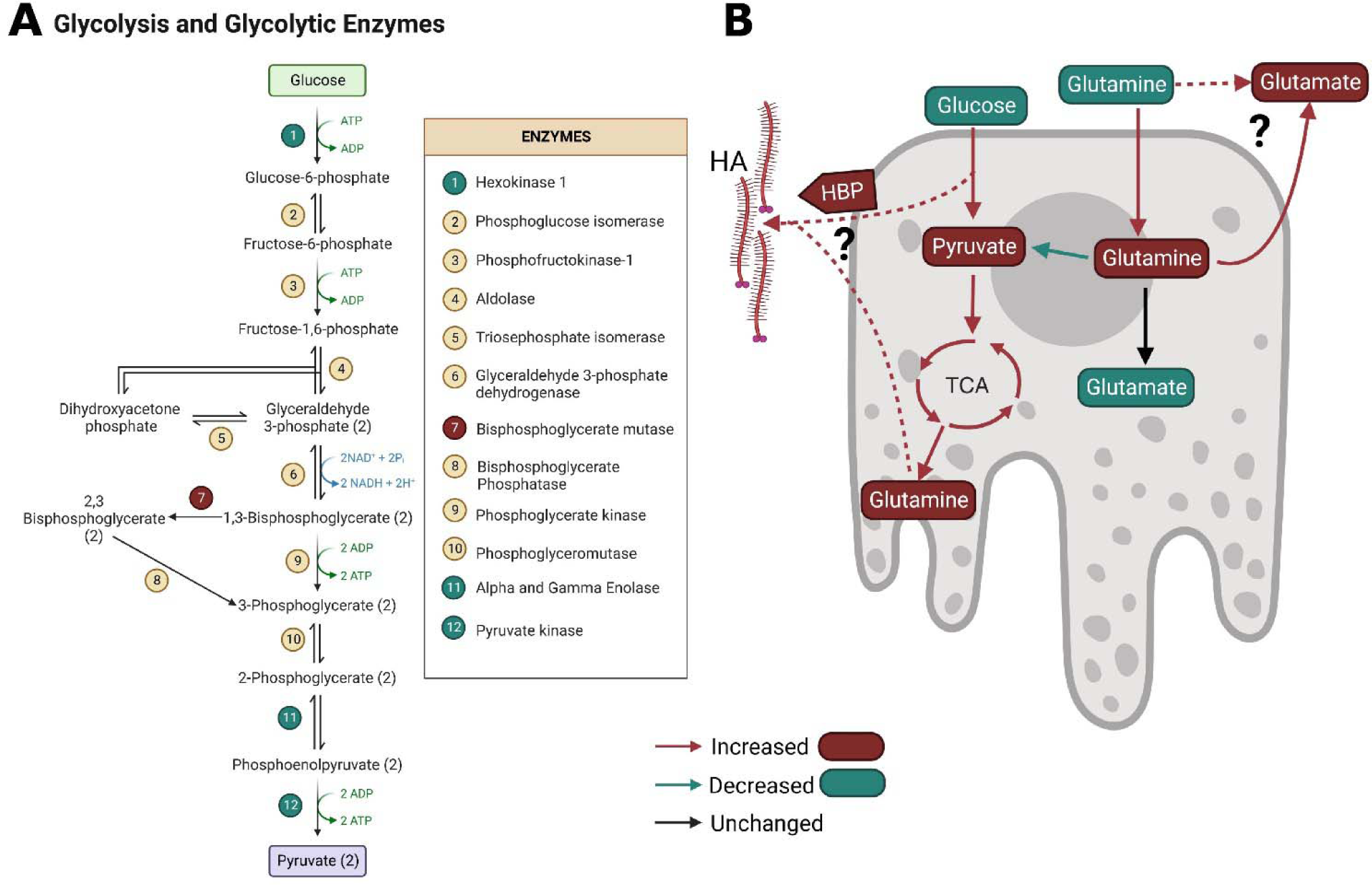
Dysregulated glycolytic enzymes and a summary of metabolic dysregulation in SFD RPE. . A, Schematic showing glycolytic intermediates and the altered glycolytic enzymes identified via proteomics. Enzymes in green circles were found to be decreased while the enzymes in red circles were found to be increased. B) Summary of metabolic dysregulation in SFD RPE. Red arrows and bubbles indicate an increase in abundance/activity while green represents a decrease in abundance/activity. Glucose utilization increases and intracellular pyruvate accumulate while glucose contribution to the TCA cycle and glutamine cataplerosis increases. Glutamine utilization increases and extracellular glutamate accumulates as intracellular glutamate decreases and intracellular glutamine accumulates. The possibility of extracellular metabolism occurring in unknown (represented by a question mark) Glutamines conversion to pyruvate via malic enzyme is decreased while glutamine anaplerosis, or contribution to the TCA cycle, is unchanged. It remains to be determined which pathway is metabolizing the additional glutamine taken up from the media and generated from glucose. We hypothesize that excess glutamine and glycolytic intermediates could be used to generate hyaluronic acid, indicated by a question mark, through the hexosamine biosynthetic pathway.

A closer look at the proteins altered in SFD RPE suggests that metabolic dysfunction spreads beyond central glucose metabolism. Panther GO pathway analysis additionally identified dysregulated proteins overrepresented in the pyrimidine, pentose phosphate and 5-hydroxytryptamine degradation pathways (**Supporting Information 1**). Proteins related to mitochondrial, fatty acid, and glutathione metabolism were also dysregulated (**Supporting Information 1**). Altered mitochondrial proteins included mitochondrial pyruvate carrier 2 (MPC2), mitochondrial glutamate carrier 1 (GC1), and ATP synthase delta and epsilon subunits suggesting mitochondrial dysfunction. Altered proteins associate with fatty acid metabolism include carnitine O-palmitoyltransferase 1, very-long-chain enoyl-CoA reductase (Tecr), and long-chain fatty acid transport protein 4 (Slc27a4). Altered glutathione-related genes includes glutathione S-transferase A4 (GSTA4), glutathione S-transferase omega-1 (GSTO1) and glutathione S-transferase Mu 2 (GSTM2), which are important for oxidative stress defense. This supports previous reports that SFD iRPE cells have impaired oxidative stress response^51,2^

While our results clearly demonstrate that TIMP3 mutations associated with SFD result in changes to RPE metabolism, the molecular mechanism by which this occurs are unclear. One possibility is that TIMP3 can directly interact with metabolic enzymes. A high throughput study identified 18 TIMP3 binding partners, one of which was ADP dependent glucose kinase which is responsible for converting glucose to glucose-6-phosphate^52^. Another possibility is that mutant TIMP3 regulates the extracellular matrix via matrix degrading enzymes which has the potential for changing cellular metabolism. Patients with SFD and AMD demonstrate significant alterations in Bruch’s membrane structure. SFD TIMP3 mutations result in highly dysregulated ECM^2^ which has the potential to alter metabolism in various ways. ECM biomechanics has previously been demonstrated to regulate intracellular metabolism in cancer^53^. Furthermore, ECM components, including HA, can act as ligands to cell surface receptors such as CD44, that trigger signaling cascades that regulate metabolism^54^. ECM constituents can also potentially be used as a nutrient source^55,56^. Exploring the relationship between ECM and metabolism could provide valuable insights into the molecular mechanisms underlying retinal degeneration.

### Experimental Procedures

#### Animals

Heterozygous *Timp3^+/S^*^179^*^C^,* a generous gift from Dr. Bernhard Weber^57^ were crossed to generate homozygous *Timp3^S^*^179^*^C^*^6^*^/S^*^179^*^C^* mice and their age-matched littermate controls. All experimental procedures were approved by the Animal Care and Use Committee (IACUC) guidelines of the Cleveland Clinic and conformed to the National Institutes of Health Guide for the Care and Use of Animals in Research and the ARVO statement for the use of animals in ophthalmic and vision research.

#### Proteomics Preparation

After euthanization, tissue was collected from 8 animals of each genotype (WT or TIMP3 S179C, 4 males and 4 females). Retina and RPE lysates were prepared from the enucleated eyes of 7 month old mice as described previously^58^. Protease buffer (200uL containing 100mM TEAB, 2% SDS, 1mM DDT) was added to each RPE/eye cup and incubated on ice for 1 hour. RPE was dissociated from choroid/sclera with vigorous tapping of the tube. RPE cells were lysed by drawing cells through a 26-27-gauge needle 15-20 times. Tissue was sonicated for 10-15 seconds at medium/frequency, centrifuged (14,000 rpm for 10 min at 4°C) and the supernatant was used for analysis.

The protein concentration of the soluble protein fraction was estimated by the Bicinchoninic Acid assay (BCA) assay. Each fraction was reduced with DTT (10mM at room temperature) alkylated with iodoacetamide (40 mM for 1h at room temperature) then quenched with DTT (40 mM)^59^. Two volumes of ice-cold acetone was used to precipitate proteins. Protein pellets were resuspended in triethylammonium bicarbonate buffer (TEAB) buffer (100mM containing 0.5 mM CaCl2) and were digested overnight at 37 °C with trypsin (initially with 2% trypsin (w/w), followed in 2 h with another 2% (w/w), and the next day with another 1% (w/w) for 2h). AccQ-Tag amino acid analysis was used to quantify soluble peptides^60,61^. Gender-specific pooled WT samples were prepared for proteomic analyses by combining equal amounts of proteolyzed protein from 3 male and 4 female WT controls.

#### iTRAQ-Labeling and Peptide Fractionation

iTRAQ-Labeling of tryptic peptides with an 8-plex iTRAQ kit was performed as previously described^61,62^ Tryptic digests of the individual KI samples (21 ug each) and the pooled reference samples (21 ug each) were each labeled with a unique iTRAQ tag and combined in a total of 2 batches as follows. Batch 1 contained 6 iTRAQ tags among the pooled male and female reference samples, 2 individual male mutant samples and 2 female mutant samples. Batch 2 was a duplicate of Batch 1 but with different KI samples. Each batch was individually fractionated by reverse-phase high performance liquid chromatography (RP-HPLC) at pH 10 using methods detailed elsewhere^62^ and chromatography fractions encompassing the entire elution were selectively combined and dried. Ten fractions per batch were analyzed by LC MS/MS.

#### Protein Identification

RP-HPLC fractions were analyzed by LC-MS/MS on an Orbitrap Fusion Lumos Tribrid mass spectrometer^61,62^. Protein identification utilized the Mascot 2.6.2 search engine and the UniProt mouse database version 2019_07 (17,026 sequences). Database search parameters were restricted to three missed tryptic cleavage sites, a precursor ion mass tolerance of 10 ppm, a fragment ion mass tolerance of 20 mmu and a false discovery rate of ≤ 1%. Protein identification required detection of a minimum of two unique peptides per protein. Fixed protein modifications included N-terminal and ε-Lys iTRAQ modifications and S-carbamidomethyl-Cys. Variable protein modifications included Met oxidation, Asn and Gln deamidation and iTRAQ Tyr. A minimum Mascot ion score of 25 was used for accepting peptide MS/MS spectra.

#### Protein Quantitation

The iTRAQ tag intensities on male and female KI mouse peptides versus their gender-specific pooled WT control were quantified by the weighted average method^63^ using the Mascot 2.6.2 Summed Intensities Program. Protein quantitation required a minimum of two unique peptides per protein, utilized reporter ion tolerance of 10 ppm, and a Mascot peptide ion score ≥ 25. Protein ratios were determined in log space and transformed back to linear ratios for reporting.

#### Statistical Analyses of the Proteomic Data

Quantile normalization was used to normalize the mass spectrometric iTRAQ proteomics data. Batch effects were examined. Significantly elevated and decreased proteins were sought using the limma package in R, and the results were adjusted for multiple-testing using the Benjamini-Hochberg procedure^57,64^. Criteria for significantly elevated and decreased proteins included average protein ratios (TIMP3 mutant/ gender specific WT) above or below the mean by at least one standard deviation with adjusted p-values ≤ 0.05. All Proteins found to be significantly altered (Adjusted P value < 0.05) between control and TIMP3 S179C were assigned a biological process or protein classification using Panther Gene Ontology v17.0 with the “gene list analysis” tool.

#### RNAseq Library preparation

Enucleated eyes from 8 total mice (11 months in age) were used: 4 TIMP3 S179C homozygous and 4 WT littermate controls containing a combination of males and females. Posterior eye cups (RPE/choroid/sclera) were isolated (as described above) and incubated with RNAprotect Tissue Reagent (Quiagen 76104) for ten minutes at room temperature with gentle agitation until the RPE was detached from the choroid and sclera. RPE cells were isolated and pelleted by centrifugation. Pellets were resuspended in QIAzol lysis buffer (Qiagen 79306) following which RNA was extracted using the RNeasy Plus Mini Kit (Qiagen 74134). The total yield of RNA was 300-900ng. The quality of the RNA was assessed by the Agilent Bioanalyzer, and each received an RNA Integrity Score of at least 8 out of 10, indicating high quality RNA was extracted. Libraries were prepared for sequencing using the TruSeq RiboZero Kit for Human/Mouse/Rat (Illumina). Ribosome-depletion was used instead of polyA selection to enrich the mRNA in the samples.

#### RNAseq and Data analysis

The libraries were sequenced on an Illumina HiSeq 2500, yielding > 30 million, 100-nucleotide paired-end reads. RNA sequencing output (fastq files) were analyzed for quality control for each sample using FastQC (https://www.bioinformatics.babraham.ac.uk/projects/fastqc/). Reads from fastq files were aligned to the genome using HISAT2 Version 2.0.5^65^. Alignment of reads to the mm9 genome was ∼93-96% in each sample. SAMtools^66^ was used to convert HISAT2 output to sorted bamfiles and to create an index for genome browser viewing. Cuffdiff Version 2.2.1^65^ was run comparing WT and TIMP3 S179C homozygous mutants using the UCSC annotation of genes in the mm9 genome. Point mutations in TIMP3 were confirmed using the Integrative Genomics Viewer (IGV)^67^. All transcripts found to be significantly altered (adjusted p value < 0.05) between control and TIMP3 S179C were assigned a biological process or protein classification using Panther Gene Ontology v17.0 with “gene list analysis” tool.

#### ARPE -19 Cell culture

Cells were cultured using protocol adapted from Hazim et al. 2019^36^. ARPE-19 were expanded using DMEM: F12, 10% FBS (Gibco #A526801, 1% Penicillin/Streptomycin until confluent within a T75 flask. After becoming confluent, cells were allowed to further differentiate in RPE media (MEM alpha with GlutaMAX (Thermo, #32561037), 1% FBS, 1% Penicillin/Streptomycin, 1% N1 supplement (Sigma, #N6530), taurine (0.25 mg/ml, Sigma #Y0625), hydrocortisone (20 ng/ml, Sigma #50396), triiodo-thyronin (0.013 ng/ml, Sigma, #T6397), and nicotinamide (10mM Sigma #N0636) which yielded a homogenous and polarize monolayer. Cells were dissociated with Trypsin/EDTA (0.025% EDTA) for 5 minutes at 37°C. The reaction was quenched with DMEM: F12, 10% FBS, 1% Penicillin/Streptomycin before being spun down and resuspended in RPE media. Cells were plated in flat bottom 6-well tissue culture plates at a seeding density of 1.66 X 10^5^ cells/cm^2^ in RPE media. Cells were maintained at 37°C and 5% CO_2._ Media was changed 2 times per week for 4 weeks until isotopic tracing experiments.

#### ARPE19 glucose isotopic tracing

To ensure that the cells had adequate nutrients before labeling started, fresh (unlabeled) cell culture media was given to the cells 24 hours prior. At the start of the experiment (time 0) unlabeled media was aspirated and wells were quickly washed with 3mL warm normal saline (0.8% NaCl). Then, 2mL of media containing [^13^C_6_] glucose was added. The media had a base of DMEM without glucose, without pyruvate, and without L-glutamine (Lerner Research Institute Central Cell Services Media Laboratory, 99AC500CUST) with 10mM [^13^C_6_] labeled glucose (Cambridge Isotope Laboratories, 99%, Cat. # CLM-1396-PK.), 2.5 mM unlabeled glutamine, 10% dialyzed FBS, 1% Penicillin/Streptomycin antibiotics. Cells were incubated with labeled media at 37°C for 24 hours.

#### ARPE-19 metabolite extraction from cells and media

After 24 hours, media was collected, flash frozen, and stored at - 80°C until analysis. Cells were rinsed with 1mL warm normal saline (0.9% NaCl, Sigma S5886, Millipore ZEQ7000T0C). Cells were immediately put on ice and 1mL of ice cold 80% Methanol (Fisher Scientific A454-4) containing 0.05 mg/ml ^13^C Ribitol internal standard was added to each well. Wells were scraped and all contents were transferred to a fresh 1.5mL tube and immediately put on wet ice. Samples were briefly sonicated to ensure lysis (∼5 seconds) and then centrifuged (15,000 x g for 5 minutes at 4°C). 500 μL of supernatant was transferred to a fresh 1.5mL tube. Samples were dried in a centrivap concentrator (Labconco 7310021) over-night at 4°C and immediately derivatized the next morning.

Media samples stored in -80°C were thawed on wet ice and vortexed. To 20 μL of media in a fresh tube, 70 μL of -20°C Methanol and 10 μL of 0.05mg/ml 13C Ribitol standard was added, vortexed, and placed on ice. Samples were centrifuged at 15,000 x g for 10 mins at 4°C. 50 μL of supernatant was moved to a fresh tube and dried overnight in Centrivap at 4°C.

#### ARPE-19 derivatization protocol

Samples were placed in Centrivap at 40°C for 5 mins to ensure removal of all moisture then immediately derivatized. 25 μL of Methoxyamine hydrochloride (Sigma 593-56-6)/pyridine (Sigma 110-86-1) solution (40 mg/ml) was added to each tube. The samples were mixed at 45°C at 1,000 rpm for 30 min on the ThermoMixer (Eppendorf 5382000023). 75 μL of MSTFA + 1% TMCS (Thermo Scientific 48915) was added to each tube. Samples were incubated at 45°C at 1,000 rpm for 30-minutes on the ThermoMixer. Samples were centrifuged for 2 minutes at room temperature at 15,000 x g. 68 μL of supernatant was put in GC/MS vials (Thomas Scientific 1152P35, Macherey-Nagel 702716).

#### ARPE-19 Mass Spectrometry/Gas Chromatography method

One microliter of each sample was injected into the 7890B GC connected to 5977 MSD Agilent GCMS system. Injections were made in splitless mode. The fused silica capillary column used was a DB-5MS 301m1×10.251mm1×10.251µm with a 10 m of built in guard column (Agilent 122-5532G). The front inlet was set to 250°C with septum purge flow of 3 ml/min of helium. Samples were analyzed in a constant flow mode with helium set to 1.1 ml/min. The GC method was 1 min at 60°C, followed by 10 °C/min increments until 325 °C and finally held at 325 °C for 10 min. Pyruvate and lactate were measured by select ion monitoring (SIM). Ions 174-177 were monitored for pyruvate from 6.6-7.7 minutes while ions 219-222 were monitored for lactate from 7.7-9 minutes. Glucose in media was measured in full scan mode using electron ionization with a scan widow from 50 to 8001m/z. A solvent delay of 6.6 minutes was applied.

#### APRE-19 Data analysis

Agilent Masshunter was used to quantify peak height for M_0_-M_n_. Relative abundance of total metabolites was calculated by adding together all isotopologues (M_0_-M*_n_*) and dividing it by the height of the internal standard, ribitol. Isocor was used to correct for natural isotopic abundances and calculate fractional enrichment. Corrected abundance output from Isocor was used to calculate the relative amount of labeled (M_0_) and unlabeled (M_1_-M*_n_*) metabolites and was normalized to ribitol internal standard.

#### Reprogramming of control and patient fibroblasts into iPSCs

iPSCs were reprogrammed from control and patient fibroblasts (confirmed for TIMP3 S204C mutation) as described previously^37,39^ Briefly, fibroblasts were electroporated with non-integrating episomal vectors containing reprogramming factors (pCXLE-hOCT4-shP53, pCXLE-hSK and pCXLE-hUL)^37,39,68^. Electroporation was carried out using the nucleofection kit for primary fibroblasts (Lonza) and Nucleofector 2b Device (Lonza, Program T-016). Following this, the electroporated cells were cultivated in fibroblast medium (DMEM with high glucose, 10% fetal bovine serum, 2 mM glutamine, and 100 U/mL penicillin-streptomycin) for 6 days. After 6 days, reprogrammed fibroblasts were trypsinized and plated onto irradiated mouse embryonic fibroblast (MEF) feeder cells and maintained in DMEM/F12 (Gibco, ThermoFisher Scientific) with 20% Knockout serum replacement (Gibco, ThermoFisher Scientific), 1% MEM-NEAA (Gibco, ThermoFisher Scientific), 1% glutamax (Gibco, ThermoFisher Scientific), and 100 ng/mL FGF2; Peprotech). iPSC colonies appeared at ∼day 17-30 from transfection and were manually collected for further expansion.

#### Differentiation of control and patient iPSCs into iRPE

iPSCs were routinely maintained on Matrigel (Corning) in mTeSRTM (Stemcell Technologies) and differentiated into iRPE using previously described protocol^37,39,69,70^. Briefly, confluent iPSC colonies were initially dissociated with ReLeSR™ and cultured as free-floating embryoid bodies (EBs) in T-25 flasks. Plating of EBs were laminin-coated 6 well plates was done on day 6 and maintained in neural induction medium (NIM: DMEM/F12 containing 1% MEM-NEAA (Gibco, ThermoFisher Scientific), 1% Glutamax (Gibco, ThermoFisher Scientific), 1% N2 Supplement (Gibco, ThermoFisher Scientific) and 2 mg/mL heparin). On day 14, the EBs were switched to retinal differentiation medium (RDM). RDM was prepared by using 70% DMEM (Gibco, ThermoFisher Scientific, #11965-092), 30% F-12 (Cytiva #SH30026.01), 2% B-27 supplement without Vitamin A (Gibco, ThermoFisher Scientific) and 1% (v/v) GibcoTM Antibiotic-Antimycotic (ThermoFisher Scientific). RPE started to appear at around day 60-90 timepoint in the differentiated iPSC cultures. Patches of RPE was dissected were dissected out under a microscope followed by dissociation with 0.05% Trypsin-EDTA (Gibco, ThermoFisher Scientific) and further plated onto laminin-coated (Gibco, ThermoFisher Scientific) 24-well plates with RDM containing 10% FBS for 2 feedings and changed to RPE containing 2% FBS until they formed a confluent monolayer and then switched to RDM only. This was considered as passage 0 (P0) RPE monolayers. Control and patient iRPE cultures were passaged as needed. Cells were only used at passage 3 and below.

#### Generation of gene corrected iRPE Control line

The day before transfection, hiPSCs were dissociated into single cells using TrypLE and seeded onto Matrigel coated 96-well plate at a density of ∼3-4 x 104 cells per well in mTeSR plus medium supplemented with 10 µM ROCK inhibitor (Y-27632; STEMCELL Technologies). Alt-R CRISPR-Cas9 crRNA(5’-TGATGATGCATTTATCCGGG-3’), tracrRNA ATTOTM 550 (Integrated DNA technologies) were resuspended and crRNA: tracrRNA duplexes were prepared according to the manufacturer’s instructions. A transfection complex was prepared with 6 nmol of Alt-R™ SpCas9 nuclease V3 (Integrated DNA technologies), 12 nmol crRNA: tracrRNA and 50 nmol of ssODN donor construct (5’GCATCCGGCAGAAGGGCGGCTACTGCAGCTGGTACCGAGGATGGGCCCCTCCGGATAAAAGCATCATCAATGCCACAGACCCCTGAGCGCCAGACCCTGCCCCACCTCAC3’) using Lipofectamine Stem Transfection Reagent (Cat. No. STEM00001; Thermo Fisher Scientific) according to manufacturer’s instructions and added dropwise to the well. The media was replaced after 24 hours and on day 3 post-transfection, cells were dissociated using TrypLE and seeded at low density (∼300-500 cells) in each well of 6-well plate). The clones were manually picked and expanded. For screening, gDNA PCR was performed using primers TIMP3-F (5’CCTGCTACTACCTGCCTTGC) and TIMP3-R (5’GGGTGGGAATTACAATCCTCAAATG) followed by restriction digestion using enzyme Nsil (New England Biolabs, Cat no. R0127L). Sanger sequencing was performed to verify the mutation correction.

#### iRPE transportation

Cells were transported by car in a portable incubator (VEVOR XHC-25) from The University of Rochester (Rochester, NY) directly to Cleveland Clinic Cole Eye Institute (Cleveland, OH). Cells were maintained at 37°C +/- 1°C during the 5 hours of transit. Media was supplemented with 25mM of HEPES buffer (Gibco #15630080) to maintain media buffering capacity without CO2 supplementation. Immediately upon arrival, media was replaced with fresh RDM media. Cells were maintained at 37°C, 5% CO_2_, 95% humidity for at least one month before experiments were conducted.

#### iRPE isotopic tracing

Forty-eight hours before experiments, fresh RDM media was added to ensure that the cells had adequate nutrients before labeling was initiated. Isotopic tracing was conducted in DMEM media without glucose, without pyruvate and without glutamine, (Cleveland Clinic Custom) supplemented with 10mM labeled or unlabeled glucose and 2.5 mM labeled or unlabeled glutamine. Glucose tracing experiments contained 10mM uniformly labeled [^13^C_6_] Glucose and 2.5 mM unlabeled glutamine. Glutamine tracing experiments contained 10mM unlabeled glucose and 2.5mM uniformly labeled [^13^C_5_] glutamine. Cells were incubated (37°C, 5% CO_2_, 95% humidity) with 1mL of glucose or glutamine labeled media for 24 hours.

#### iRPE metabolite extraction

After 24 hours, the media was collected, and flash frozen for extracellular metabolomics. Cells were quickly washed once with 1mL of chilled normal saline (0.9% NaCl). Intracellular metabolism was quenched, and metabolites extracted by adding 200 μL of ice cold 80% methanol containing 13C ribitol and Norvaline (Waters 186009301) as internal standards. Plates were immediately put on ice and cells were detached using plastic cell scraper. To ensure full cell lysis, cells were sonicated at 4°C for 30-40 seconds. Proteins were precipitated out through centrifugation at 15,000 x g for 5 minutes at 4°C. 25 μL of supernatant was transferred to a fresh 1.5 mL tube before being dried at room temperature using a SpeedVac concentrator (Savant SC110-120). Samples subsequently were stored at -80°C.

Extracellular media samples were thawed on ice and metabolites were extracted from 5µL of media with 95µL extraction buffer (ice cold 80% methanol containing 5 μL of 0.05mg/ml ribitol and 5 μL of 5.86pg/µL Norvaline as internal standards) Samples were centrifuged at 15,000 x g for 10 mins at 4°C. 50 μL of supernatant was moved to a fresh tube and dried at room temperature for 20 min with Savant SpeedVac concentrator.

#### iRPE derivatization

Derivatization protocol was adapted from previously published protocols^71^. Briefly, to ensure samples were completely dry, they were put into the SpeedVac concentrator for 5 minutes. 10μL Methoxyamine/pyridine (40mg/1mL) was added to dry samples. Using an Eppendorf ThermoMixer® C, samples were mixed at 300 rpms for 1.5 hours at 37°C. 30 μL MTBSTFA (N-tert-Butyldimethylsilyl-N-methyltrifluoroacetamide, MilliPore Sigma, 394882) was added and mixed on the thermomixer at 300rpms for 30 minutes at 70°C. Samples were centrifuged for 10 minutes at room temperature before 30 μL of supernatant was moved to Crimp Vials with glass inserts and capped.

#### GC/MS analysis

Methods were adapted from previous publications^71^. An auto sampler (Agilent 7693A ASL) was used to inject 1 μL in splitless mode using a 5 μL syringe (Agilent G4153-80213) into an Agilent 8890 GC followed by an Agilent 5977C MSD. Inlet was lined with a splitless fritted liner (Agilent 5190-5112-025) at a temperature of 250°C and pressure of 14.067 psi. To avoid carry-over, the syringe was washed twice before injection and three times after injection. Each wash consisted of 2µL hexane followed by 2µL acetone. The total inlet flow was 54 mL/min and septum purge flow was 3mL/mins in standard mode. Purge flow to split vent was 50 mL/min at 0.5 min. The column used was an ultra-inert DB-5ms (UI) with 30m length x 0.25mm inner diameter x 0.25 µm film thickness (Agilent 122-5532UI) with a constant flow of 1mL/min. Initial oven temperature was 95°C with a 2-minute hold time and was ramped to 270°C at a rate of 10°C/minute. Oven was ramped a second time at 5°C/minute until 300°C was reached and held for 6 minutes. MSD transfer line was constantly kept at 290°C. MSD was run full scan mode (50-600m/z) with a solvent delay of 5.95 minutes. MSD Source and quad were set to 250°C and 150°C, respectively.

#### Quality Control

Each run contained 2 solvent blanks to assess carry-over, 2 derivatization blanks to access background, and 2 biological quality control samples made from pooled porcine RPE that were extracted the same way as experimental samples. After derivatization, biological QC samples were pooled once again to ensure uniformity before being separated into two separate vials for GC/MS analysis. One QC samples was injected 3 times at the beginning of the sequence and the other QC sample was injected 3 times at the end to confirm reproducibility and evaluate machine drift over the run. Each metabolite was evaluated separately for reducibility by quantitating the area under the curve (AUC) normalized to total sum. Only metabolites that had a relative standard deviation under 7% across all six QC injections were included in results.

#### Data Analysis and statistics

The area under the curve was quantified with Agilent MassHunter Quantitative Analysis. Compounds, retention times, and quantitative m/z can be found in **Supplemental methods 1.** Total relative metabolite abundance was calculated by summing the area together of all isotopologues (M0-M*_n_*) normalized with total sum of measured metabolites (mTIC). Fold change relative to Crispr corrected controls was calculated by taking the normalized metabolite level divided by the average of the control samples. Multiple unpaired t. tests were performed using GraphPad Prism version 10.0.0 for Windows, GraphPad Software, Boston, Massachusetts USA, www.graphpad.com

## Data Availability

RNAseq data will be uploaded to Gene Expression Omnibus (https://www.ncbi.nlm.nih.gov/geo/), Proteomics data will be uploaded to Proteomics Identifications Database (PRIDE Archive, https://www.ebi.ac.uk/pride/archive/), and Metabolomics data will be uploaded to Metabolomics Workbench (https://www.metabolomicsworkbench.org/). Additional Data is available upon request.

## Supporting Information

**Supporting Information 1:** Proteomics Results including all the significantly different proteins (adjusted p < 0.05) identified in males (tab titled “Males”), females (“Females”), and the proteins altered in both males and females (“Male_female_common”). Panther GO results from biological process (“biological process”), protein class (“protein class”), and pathway analysis (“pathway”). List of significantly different proteins corresponding with metabolic process (“metabolic process”), metabolite interconversion (“metabolite interconversion”), metabolic pathways (“pathway genes”). This also includes a tab that has a list of all the UniProt identifiers of the significantly altered genes used for Panther Go analysis (“all_sig”).

**Supporting information 2:** Significantly altered transcripts identified via RNAseq used for Panther GO analysis (“Significant Transcripts”). Panther GO results from biological process (“Biological process”), protein class (“protein class”), and pathway analysis (“pathway”). List of significantly different transcripts corresponding with metabolic process (“metabolic process”), metabolite interconversion (“metabolite interconversion”).

**Supporting information 3:** Comparison of the cellular processes and protein classes identified by proteomics and RNAseq and genes found in both. Sheets include the number of differential genes identified by Panther GO for each category. Also include genes that were dysregulated in both data set (“overlapping genes”).

**Supporting information 4:** Full mass isotopic distributions for [^13^C_6_] Glucose isotopic tracing

**Supporting information 5:** Full mass isotopic distributions for [^13^C_5_] Glutamine tracing

**Supplemental Methods 1:** iRPE Mass spec/data analysis method and standards used to confirm peak assignments.

## Acknowledgments

The authors thank Dr. Henri Brunengraber and the Mouse Metabolic Phenotyping Center for guidance on isotopic tracing experiments, Noah Chernosky for his assistance with transporting iRPE cells, and the laboratory of Dr. Jonathan Sears for sharing supplies and analytical standards for GC/MS analysis. OpenAI’s ChatGPT3 was utilized for rephrasing to improve clarity and checking grammar. Figures were made using BioRender, Graphpad Prism, and Adobe Illustrator

## Funding and additional information

This research was supported by NIH 5 T32 EY 7157-19, R01 EY 027083, P30 EY 025585, 1F31EY033223-01, RPB Center Grant, Cleveland Eye Bank Foundation, and Cleveland Clinic Foundation support.

## Figures

**Figure S1:**
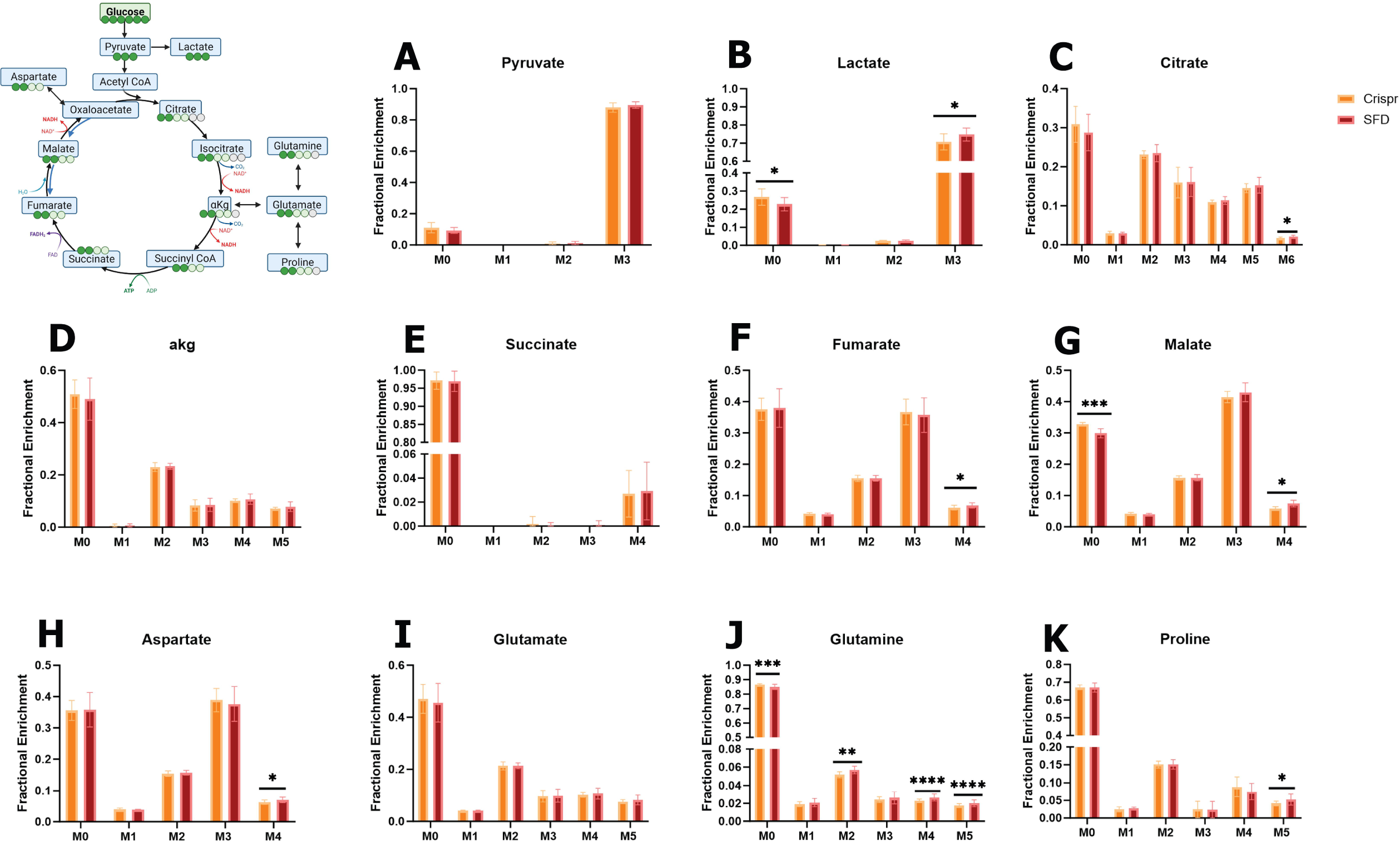
Mass isotopic distrubtions from 13C6 Glucose labeling of SFD iRPE (SFD, red) com­ paared with crispr corrected controls (Crispr, oranage)Statistics performed were 2 tailed unpaired T.tests. *p > 0.05, **p > 0.01, *** p > 0.001, **** p > 0.0001. Experimental schematic was generated using BioRender. Graphs were made with GraphPad Prism

**Figure S2:**
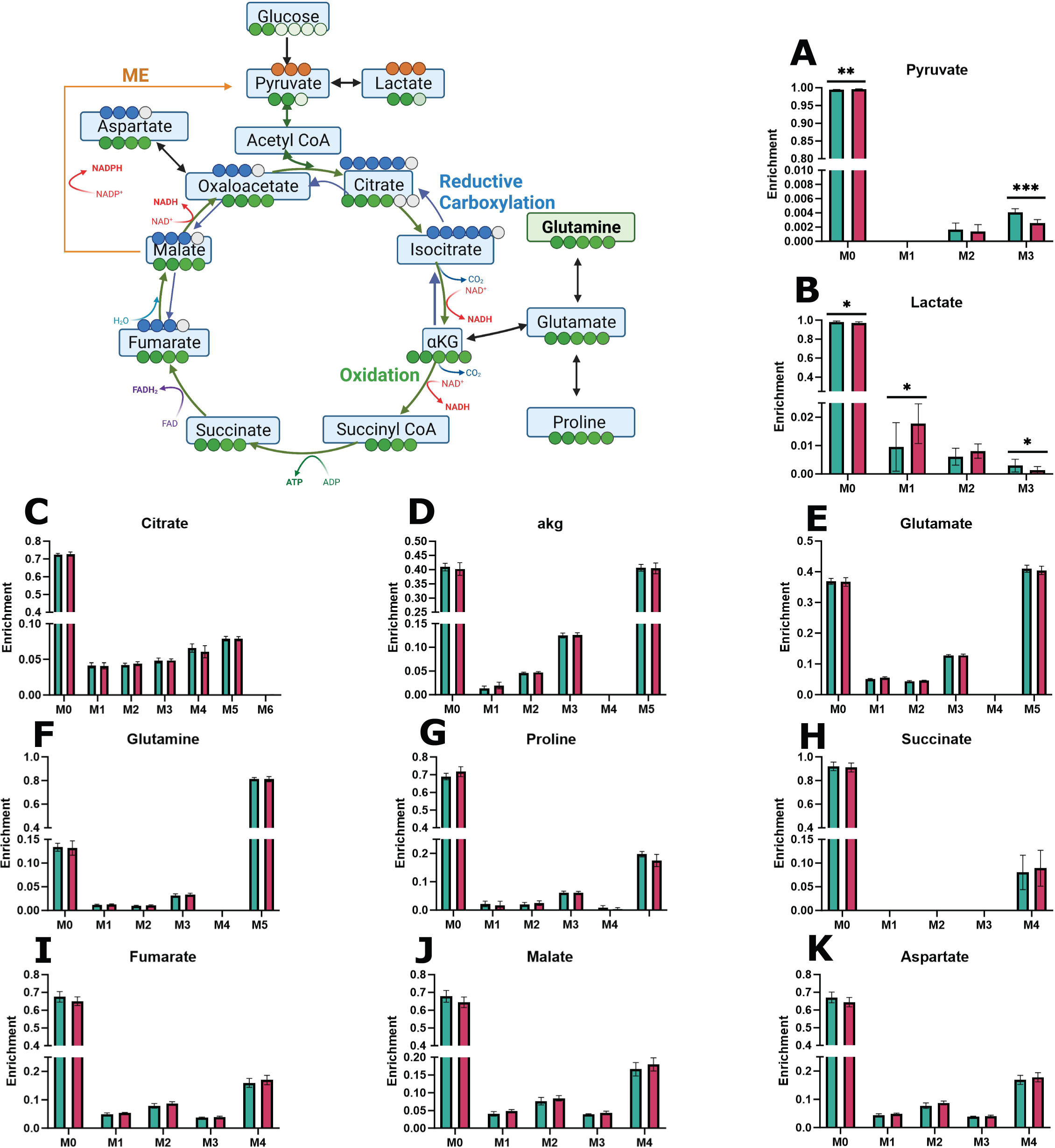
Mass isotopic distrubtions from 13C5 glutamine labeling of SFD iRPE (SFD, pink) com­ paared with crispr corrected controls (Crispr, green)Statistics performed were 2 tailed unpaired T.tests. *p > 0.05, **p > 0.01, *** p > 0.001, **** p > 0.0001. Experimental schematic was generated using BioRender. Graphs were made with GraphPad Prism

## References

1. Capon, M. R. et al. Sorsby’s fundus dystrophy. A light and electron microscopic study. Ophthalmology 96, 1769– 1777 (1989).

2. Engel, A. L. et al. Extracellular matrix dysfunction in Sorsby patient-derived retinal pigment epithelium. Exp. Eye Res. 215, 108899 (2022).

3. Gourier, H. C. Y. & Chong, N. V. Can Novel Treatment of Age-Related Macular Degeneration Be Developed by Better Understanding of Sorsby’s Fundus Dystrophy. J. Clin. Med. 4, 874–883 (2015).

4. Hollyfield, J. G., Salomon, R. G. & Crabb, J. W. Proteomic approaches to understanding age-related macular degeneration. Adv. Exp. Med. Biol. 533, 83–89 (2003).

5. Fariss, R., Apte, S., Luthert, P., Bird, A. & Milam, A. Accumulation of tissue inhibitor of metalloproteinases-3 in human eyes with Sorsby’s fundus dystrophy or retinitis pigmentosa. Br. J. Ophthalmol. 82, 1329–1334 (1998).

6. Zhang, M. et al. Dysregulated metabolic pathways in age-related macular degeneration. Sci. Rep. 10, 2464 (2020).

7. Hou, X.-W., Wang, Y. & Pan, C.-W. Metabolomics in Age-Related Macular Degeneration: A Systematic Review. Invest. Ophthalmol. Vis. Sci. 61, (2020).

8. Fisher, C. R. & Ferrington, D. A. Perspective on AMD Pathobiology: A Bioenergetic Crisis in the RPE. Invest. Ophthalmol. Vis. Sci. 59, AMD41–AMD47 (2018).

9. Wang, W. et al. Metabolic Deregulation of the Blood-Outer Retinal Barrier in Retinitis Pigmentosa. Cell Rep. 28, 1323–1334.e4 (2019).

10. Nguyen, T. T. & Wong, T. Y. Retinal vascular manifestations of metabolic disorders. Trends Endocrinol. Metab. TEM 17, 262–268 (2006).

11. Abcouwer, S. F. & Gardner, T. W. Diabetic retinopathy: loss of neuroretinal adaptation to the diabetic metabolic environment. Ann. N. Y. Acad. Sci. 1311, 174–190 (2014).

12. Singh, C. Metabolism and Vascular Retinopathies: Current Perspectives and Future Directions. Diagnostics 12, 903 (2022).

13. Singh, C. et al. Serine and 1-carbon metabolism are required for HIF-mediated protection against retinopathy of prematurity. JCI Insight 4, e129398, 129398 (2019).

14. Scerri, T. S. et al. Genome-wide analyses identify common variants associated with macular telangiectasia type 2. Nat. Genet. 49, 559–567 (2017).

15. Gantner, M. L. et al. Serine and Lipid Metabolism in Macular Disease and Peripheral Neuropathy. N. Engl. J. Med. 381, 1422–1433 (2019).

16. Bonelli, R. et al. Genetic disruption of serine biosynthesis is a key driver of macular telangiectasia type 2 aetiology and progression. Genome Med. 13, 39 (2021).

17. Wang, F. et al. A Missense Mutation in HK1 Leads to Autosomal Dominant Retinitis Pigmentosa. Invest. Ophthalmol. Vis. Sci. 55, 7159–7164 (2014).

18. Zhang, L. et al. Whole-exome sequencing revealed HKDC1 as a candidate gene associated with autosomal-recessive retinitis pigmentosa. Hum. Mol. Genet. 27, 4157–4168 (2018).

19. Chen, C. T., Shao, Z. & Fu, Z. Dysfunctional peroxisomal lipid metabolisms and their ocular manifestations. Front. Cell Dev. Biol. 10, 982564 (2022).

20. Chen, Y. et al. Metabolism Dysregulation in Retinal Diseases and Related Therapies. Antioxidants 11, (2022).

21. Pan, W. W., Wubben, T. J. & Besirli, C. G. Photoreceptor metabolic reprogramming: current understanding and therapeutic implications. *Commun*. Biol. 4, 245 (2021).

22. Kurihara, T. et al. Hypoxia-induced metabolic stress in retinal pigment epithelial cells is sufficient to induce photoreceptor degeneration. eLife 5,.

23. Kumagai, A. K. Glucose transport in brain and retina: implications in the management and complications of diabetes. Diabetes Metab. Res. Rev. 15, 261–273 (1999).

24. Tang, J., Zhu, X. W., Lust, W. D. & Kern, T. S. Retina accumulates more glucose than does the embryologically similar cerebral cortex in diabetic rats. Diabetologia 43, 1417–1423 (2000).

25. Chao, J. R. et al. Human retinal pigment epithelial cells prefer proline as a nutrient and transport metabolic intermediates to the retinal side. J. Biol. Chem. 292, 12895–12905 (2017).

26. Yam, M. et al. Proline mediates metabolic communication between retinal pigment epithelial cells and the retina. J. Biol. Chem. 294, 10278–10289 (2019).

27. Adijanto, J. et al. The Retinal Pigment Epithelium Utilizes Fatty Acids for Ketogenesis. J. Biol. Chem. 289, 20570– 20582 (2014).

28. Du, J. et al. Reductive carboxylation is a major metabolic pathway in the retinal pigment epithelium. Proc. Natl. Acad. Sci. 113, 14710–14715 (2016).

29. Kanow, M. A. et al. Biochemical adaptations of the retina and retinal pigment epithelium support a metabolic ecosystem in the vertebrate eye. eLife 6, (2017).

30. Ferrington, D. A. et al. Altered bioenergetics and enhanced resistance to oxidative stress in human retinal pigment epithelial cells from donors with age-related macular degeneration. Redox Biol. 13, 255–265 (2017).

31. Oslund, R. C. et al. Bisphosphoglycerate mutase controls serine pathway flux via 3-phosphoglycerate. Nat. Chem. Biol. 13, 1081–1087 (2017).

32. Sethi, J. K. & Vidal-Puig, A. Wnt signalling and the control of cellular metabolism. Biochem. J. 427, 1 (2010).

33. Ata, R. & Antonescu, C. N. Integrins and Cell Metabolism: An Intimate Relationship Impacting Cancer. Int. J. Mol. Sci. 18, 189 (2017).

34. Dunn, K. C., Aotaki-Keen, A. E., Putkey, F. R. & Hjelmeland, L. M. ARPE-19, a human retinal pigment epithelial cell line with differentiated properties. Exp. Eye Res. 62, 155–169 (1996).

35. Qi, J. H. et al. Expression of Sorsby’s fundus dystrophy mutations in human retinal pigment epithelial cells reduces matrix metalloproteinase inhibition and may promote angiogenesis. J. Biol. Chem. 277, 13394–13400 (2002).

36. Hazim, R. A., Volland, S., Yen, A., Burgess, B. L. & Williams, D. S. Rapid differentiation of the human RPE cell line, ARPE-19, induced by nicotinamide. Exp. Eye Res. 179, 18–24 (2019).

37. Galloway, C. A. et al. Drusen in patient-derived hiPSC-RPE models of macular dystrophies. Proc. Natl. Acad. Sci. U. S. A. 114, E8214–E8223 (2017).

38. Singh, R. et al. Functional Analysis of Serially Expanded Human iPS Cell-Derived RPE Cultures. Invest. Ophthalmol. Vis. Sci. 54, 6767–6778 (2013).

39. Manian, K. V. et al. 3D iPSC modelling of retinal pigment epithelium-choriocapillaris complex identifies factors involved in the pathology of macular degeneration. Cell Stem Cell 28, 846–862.e8 (2021).

40. Grenell, A. et al. Loss of MPC1 reprograms retinal metabolism to impair visual function. Proc. Natl. Acad. Sci. U. S. A. 116, 3530–3535 (2019).

41. Tanner, L. B. et al. Four Key Steps Control Glycolytic Flux in Mammalian Cells. Cell Syst. 7, 49–62.e8 (2018).

42. Wierenga, A. T. J. et al. HIF1/2-exerted control over glycolytic gene expression is not functionally relevant for glycolysis in human leukemic stem/progenitor cells. Cancer Metab. 7, 11 (2019).

43. Henderson, A. R. BIOCHEMISTRY OF HYPOXIA: CURRENT CONCEPTS I: AN INTRODUCTION TO BIOCHEMICAL PATHWAYS AND THEIR CONTROL. BJA Br. J. Anaesth. 41, 245–250 (1969).

44. Loo, J. M. et al. Extracellular Metabolic Energetics Can Promote Cancer Progression. Cell 160, 393–406 (2015).

45. Yegutkin, G. G. Enzymes involved in metabolism of extracellular nucleotides and nucleosides: Functional implications and measurement of activities. Crit. Rev. Biochem. Mol. Biol. 49, 473–497 (2014).

46. Angulo, M. A., Royo, F. & Falcón-Pérez, J. M. Metabolic Nano-Machines: Extracellular Vesicles Containing Active Enzymes and Their Contribution to Liver Diseases. Curr. Pathobiol. Rep. 7, 119–127 (2019).

47. Göran Ronquist, K. Extracellular vesicles and energy metabolism. Clin. Chim. Acta 488, 116–121 (2019).

48. Flores-Bellver, M. et al. Extracellular vesicles released by human retinal pigment epithelium mediate increased polarised secretion of drusen proteins in response to AMD stressors. J. Extracell. Vesicles 10, e12165 (2021).

49. Wang, A. L. et al. Autophagy, exosomes and drusen formation in age-related macular degeneration. Autophagy 5, 563–564 (2009).

50. Wolk, A. et al. Role of FGF and Hyaluronan in Choroidal Neovascularization in Sorsby Fundus Dystrophy. Cells 9, 608 (2020).

51. Wolk, A. et al. The Retinal Pigment Epithelium has increased oxidative stress and is more susceptible to oxidative stress-induced degeneration in Sorsby Fundus Dystrophy. Invest. Ophthalmol. Vis. Sci. 61, 4163–4163 (2020).

52. Huttlin, E. L. et al. Architecture of the human interactome defines protein communities and disease networks. Nature 545, 505–509 (2017).

53. Deville, S. S. & Cordes, N. The Extracellular, Cellular, and Nuclear Stiffness, a Trinity in the Cancer Resistome—A Review. Front. Oncol. 9, (2019).

54. Weng, X., Maxwell-Warburton, S., Hasib, A., Ma, L. & Kang, L. The membrane receptor CD44: novel insights into metabolism. Trends Endocrinol. Metab. 33, 318–332 (2022).

55. Muranen, T. et al. Starved epithelial cells uptake extracellular matrix for survival. Nat. Commun. 8, 13989 (2017).

56. Rainero, E. Extracellular matrix internalization links nutrient signalling to invasive migration. Int. J. Exp. Pathol. 99, 4–9 (2018).

57. Weber, B. H. F. et al. A mouse model for Sorsby fundus dystrophy. Invest. Ophthalmol. Vis. Sci. 43, 2732–2740 (2002).

58. Wei, H., Xun, Z., Granado, H., Wu, A. & Handa, J. T. An Easy, Rapid Method to Isolate RPE Cell Protein from the Mouse Eye. Exp. Eye Res. 145, 450–455 (2016).

59. Ng, K.-P. et al. Retinal Pigment Epithelium Lipofuscin Proteomics. Mol. Cell. Proteomics MCP 7, 1397–1405 (2008).

60. Crabb, J. W., West, K. A., Dodson, W. S. & Hulmes, J. D. Amino Acid Analysis. Curr. Protoc. Protein Sci. 7, 11.9.1–11.9.42 (1997).

61. Saikia, P. et al. Quantitative proteomic comparison of myofibroblasts derived from bone marrow and cornea. Sci. Rep. 10, 16717 (2020).

62. Jang, G.-F. et al. Proteomics of Primary Uveal Melanoma: Insights into Metastasis and Protein Biomarkers. Cancers 13, 3520 (2021).

63. Hultin-Rosenberg, L., Forshed, J., Branca, R. M. M., Lehtiö, J. & Johansson, H. J. Defining, Comparing, and Improving iTRAQ Quantification in Mass Spectrometry Proteomics Data*. Mol. Cell. Proteomics 12, 2021–2031 (2013).

64. R-Development-Team R: A language and environment for statistical computing., R version 3.6.3, February 29, 2020; Vienna, Austria, 2019.

65. Pertea, M., Kim, D., Pertea, G. M., Leek, J. T. & Salzberg, S. L. Transcript-level expression analysis of RNA-seq experiments with HISAT, StringTie and Ballgown. Nat. Protoc. 11, 1650–1667 (2016).

66. Twelve years of SAMtools and BCFtools | GigaScience | Oxford Academic. https://academic.oup.com/gigascience/article/10/2/giab008/6137722.

67. Trapnell, C. et al. Differential gene and transcript expression analysis of RNA-seq experiments with TopHat and Cufflinks. Nat. Protoc. 7, 562–578 (2012).

68. Crombie, D. E. et al. Friedreich’s ataxia induced pluripotent stem cell-derived cardiomyocytes display electrophysiological abnormalities and calcium handling deficiency. Aging 9, 1440–1449 (2017).

69. Tang, C. et al. A human model of Batten disease shows role of CLN3 in phagocytosis at the photoreceptor–RPE interface. *Commun*. Biol. 4, 1–18 (2021).

70. Dalvi, S. et al. Environmental stress impairs photoreceptor outer segment (POS) phagocytosis and degradation and induces autofluorescent material accumulation in hiPSC-RPE cells. Cell Death Discov. 5, 1–16 (2019).

71. Xu, R., Wang, Y. & Du, J. Tracing Nitrogen Metabolism in Mouse Tissues with Gas Chromatography-Mass Spectrometry. Bio-Protoc. 11, e3925 (2021).

